# A mouse-adapted SARS-CoV-2 strain replicating in standard laboratory mice

**DOI:** 10.1101/2021.07.10.451880

**Authors:** Xavier Montagutelli, Matthieu Prot, Grégory Jouvion, Laurine Levillayer, Laurine Conquet, Edouard Reyes-Gomez, Flora Donati, Melanie Albert, Sylvie van der Werf, Jean Jaubert, Etienne Simon-Lorière

**Affiliations:** Mouse Genetics Laboratory, Institut Pasteur, Paris, France; Evolutionary Genomics of RNA Viruses, Institut Pasteur, Paris, France; Ecole Nationale Vétérinaire d’Alfort, Unité d’Histologie et d’Anatomie Pathologique, Maisons-Alfort, F-94700, France; Ecole Nationale Vétérinaire d’Alfort, BioPôle Alfort, Laboratoire d’anatomo-cytopathologie, Maisons-Alfort, F-94700, France; Univ Paris Est Créteil, EnvA, ANSES, Unité DYNAMIC, Créteil, France; Functional Genetics of Infectious Diseases, Institut Pasteur, Paris, France; Ecole Nationale Vétérinaire d’Alfort, IMRB, Inserm, F-94700 Maisons-Alfort, France; Univ Paris Est Créteil, INSERM, IMRB, F-94010 Créteil, France; Molecular Genetics of RNA viruses, CNRS UMR 3569, Université de Paris, Institut Pasteur, Paris, France; National Reference Center for Respiratory Viruses, Institut Pasteur, Paris, France

**Keywords:** SARS-CoV-2, mouse model

## Abstract

SARS-CoV-2 has infected almost 200 million humans and caused over 4 million deaths worldwide. Evaluating countermeasures and improving our understanding of COVID-19 pathophysiology require access to animal models that replicate the hallmarks of human disease. Mouse infection with SARS-CoV-2 is limited by poor affinity between the virus spike protein and its cellular receptor ACE2. We have developed by serial passages the MACo3 virus strain which efficiently replicates in the lungs of standard mouse strains and induces age-dependent lung lesions. Compared to other mouse-adapted strains and severe mouse models, infection with MACo3 results in mild to moderate disease and will be useful to investigate the role of host genetics and other factors modulating COVID-19 severity.

## Main text

Since its emergence in 2019, severe acute respiratory syndrome coronavirus 2 (SARS-CoV-2) has infected almost 200 million humans and caused over 4 million deaths worldwide, resulting also in economic crisis. To help developing and assessing preventive and therapeutic interventions, animal models have been produced in several species. Unlike other species such as non-human primate, hamsters, ferrets, minks and cats, mice show low permissiveness to SARS-CoV-2 replication due to poor binding of the virus spike protein on the rodent version of the cellular receptor angiotensin-converting enzyme 2 (ACE2) [1]. This natural resistance can be overcome either by expressing the human ACE2 receptor by transgenesis [2,3] or by transduction [4-7], or by mutating the virus in the receptor binding domain (RBD) of the spike protein to improve its binding to mouse ACE2 [8-12]. These strategies have yielded models with diverse clinical presentations. Upon SARS-CoV-2 infection, transgenic K18-hACE2 mice develop severe symptoms characterized by rapid body weight loss and death in 5-8 days [2,3]. Transduction of hACE2 in the respiratory tract confers permissiveness to viral replication to high titers but infection remains essentially asymptomatic with mild and transient body weight loss [5,6]. Mouse-adapted viral strains have been produced either by reverse genetics to modify specific residues of the RBD [8], by serial passaging in vitro or in vivo to select variants with improved replication performances [9,11,12], or by a combination of both [10]. Infection with these variants, which carry different sets of mutations, results in a broad range of models from mildly symptomatic to severe with lethality, depending on the genetics and age of mice.

### Serial passaging to adapt the virus

In an attempt to infect mice with SARS-CoV-2, we inoculated intranasally one of the first viral isolates (IDF0372) to young adult mice of C57BL/6 and of three strains of the Collaborative Cross (CC), a collection of inbred strains with large genetic diversity [13]. While none of the C57BL/6 and CC002 mice showed significant viral loads in their lungs 4 days post-infection (dpi), one in four CC001 and one in four CC071 mice had viral loads above the limit of detection (LOD), and two in three CC002 and two in four CC071 mice had detectable viral RNA (yet below 100 eqPFU/g tissue). Histological analysis of the lungs was normal in C57BL/6 mice, while the virus-positive CC mice showed peribronchiolar inflammatory infiltrates with epithelial erosion (Fig S1). This observation opened a way to attempt adapting SARS-CoV-2 to mice using young adults of CC strains. We proceeded by serial passaging first on CC071 alone and later in combination with CC001 and C57BL/6 genetic backgrounds (Fig 1A). After an initial increase, lung viral load stabilized in later generations (Fig 1B). Of note, since mice were inoculated with fresh lung homogenates before viral load was quantified, the inoculation dose was variable between passages. After 15 and 16 passages, two viral variants named MACo1 and MACo2 were isolated. Sequencing revealed a shared mutation Q493R (A22040G) located in the RBD region that directly interfaces with the receptor (Fig 1C). MACo2 was used for further passaging on BALB/c mice with the aim of favoring variants which would more efficiently replicate in standard laboratory strains. After 11 passages, another variant was isolated, cloned and named MACo3. Sequencing revealed the presence of a second mutation in the RBD: Q498R (A23055G). Two other amino-acid substitutions and one small deletion were also identified (Fig 1C). Mutations at residues 493 and 498 have been reported in other mouse-adapted strains (Fig 1D), suggesting that these positions can modulate efficient interaction between viral spike and mouse ACE2 (mACE2). MACo3 has retained the capacity to infect VeroE6 cells (suggesting preserved interaction with hACE2) and can also infect mouse cells expressing mACE, unlike the original virus (Fig 1E).

**Figure 1:**
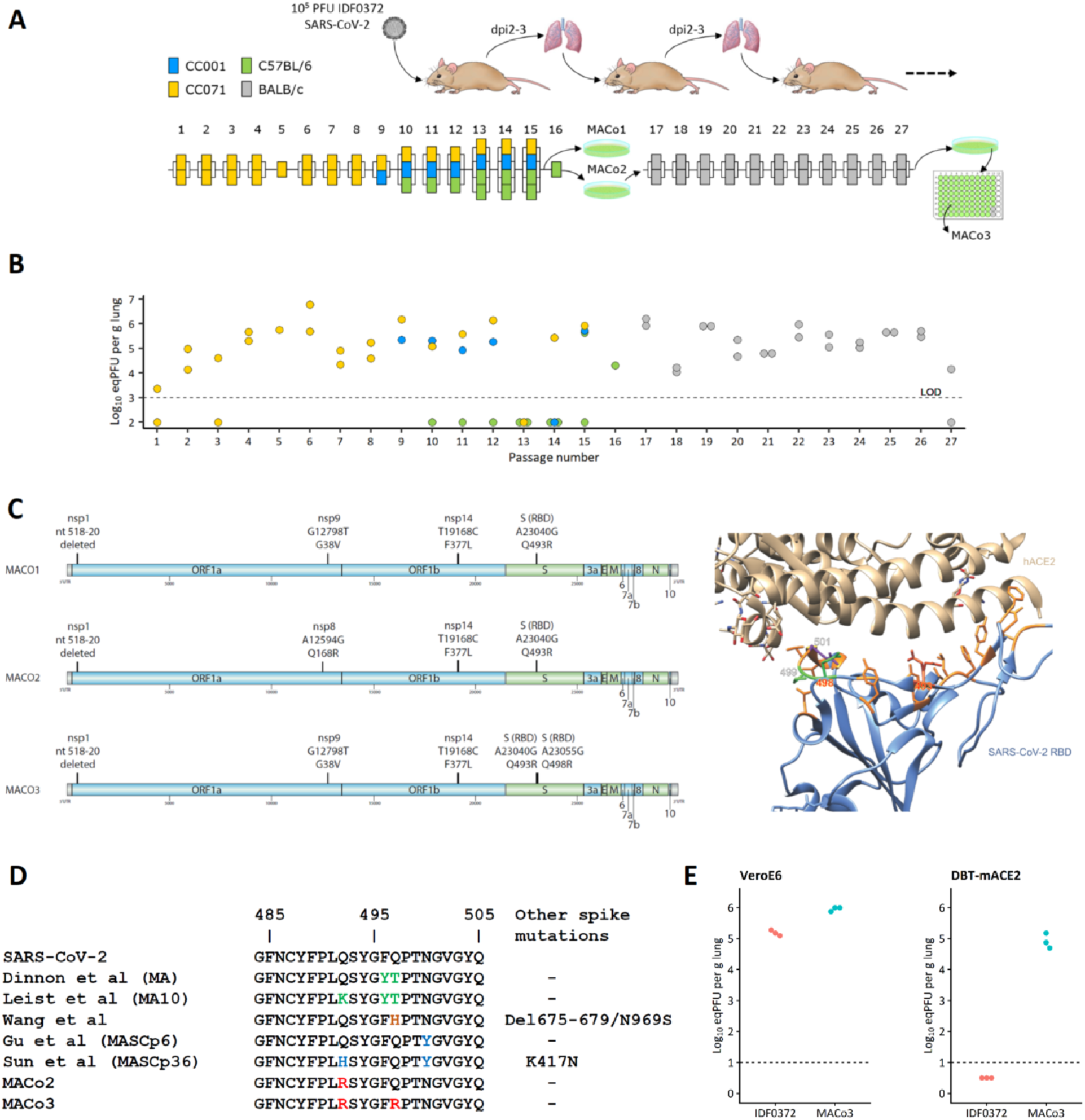
Production and characterization of the MACo3 SARS-CoV-2 strain. A: Two CC071 mice received an intranasal inoculation of 10^5^ PFU of the SARS-CoV-2 IDF0372 isolate. Their lungs were collected at dpi 3 and individual lung homogenates were pooled and used to infect mice of the next passage. Further passages involved CC071, CC001 and C57BL/6 mice until the MACo1 and MACo2 variants were isolated and characterized. Starting with MACo2, 11 more passages were performed on BALB/c mice and led to the isolation of another variant which was cloned and named MACo3. B: Viral load in individual lung homogenates increased after the first passages but remained stable later (note that the inoculation dose was variable between passages since mice were inoculated before the viral load of lung homogenates was quantified). C: Left, sequence variants found in the three MACo viruses, aligned to the proteins encoded by the viral genome. Right: Visualization of the interface between the spike RBD and the human ACE2 receptor, showing the position of the two amino acid mutated in MACo3. D: Left, alignment of the MACo1-3 variants with other mouse-adapted strains previously published. Right, viral titer in the supernatant of VeroE6 and DBT-mACE2 cells 48h after infection with IDF0372 or MACo3 viruses at an MOI of 0.1.

### Infection of standard laboratory strains with MACo3

To assess the potential of MACo3 to induce morbidity and pathological features characteristic of COVID-19 in standard laboratory mice, we inoculated intranasally young (9 week-old) and aged (12-18 month-old) BALB/c and C57BL/6 mice with 10^5^ inoculum. In young BALB/c mice, inoculation with MACo3 led to asymptomatic infection with a transient reduction of body weight (Fig 2A) and to moderate viral loads in the lung which decreased by two log10 units between dpi 3 and 6 (Fig 2B). In contrast, infection of one year-old BALB/c mice led to a rapid and durable body weight loss (peaking at 12% on dpi 3-4, Fig 2C), with mice showing symptoms of ruffled fur, hunched back posture, reduced mobility and breathing difficulties that peaked at dpi 3-6 and slowly resolved until dpi 12. Consistently, higher titers of virus were detected in the lungs of aged mice at both day 3 and 6 post inoculation (Fig 2D), compatible with slower resolution of the infection. We did not detect significant viral dissemination to other organs (Fig S2). Histopathological analysis of the lung revealed minimal broncho-interstitial pneumonia in young mice (Fig 2E), and more severe lesions (mild to moderate) in aged mice (Fig 2F). Alterations included interstitial inflammation, sometimes centered on blood vessels (perivasculitis), alteration and necrosis of bronchial and bronchiolar epithelial cells, and, in aged mice especially, endothelial cell injury and inflammation (endothelitis).

**Figure 2:**
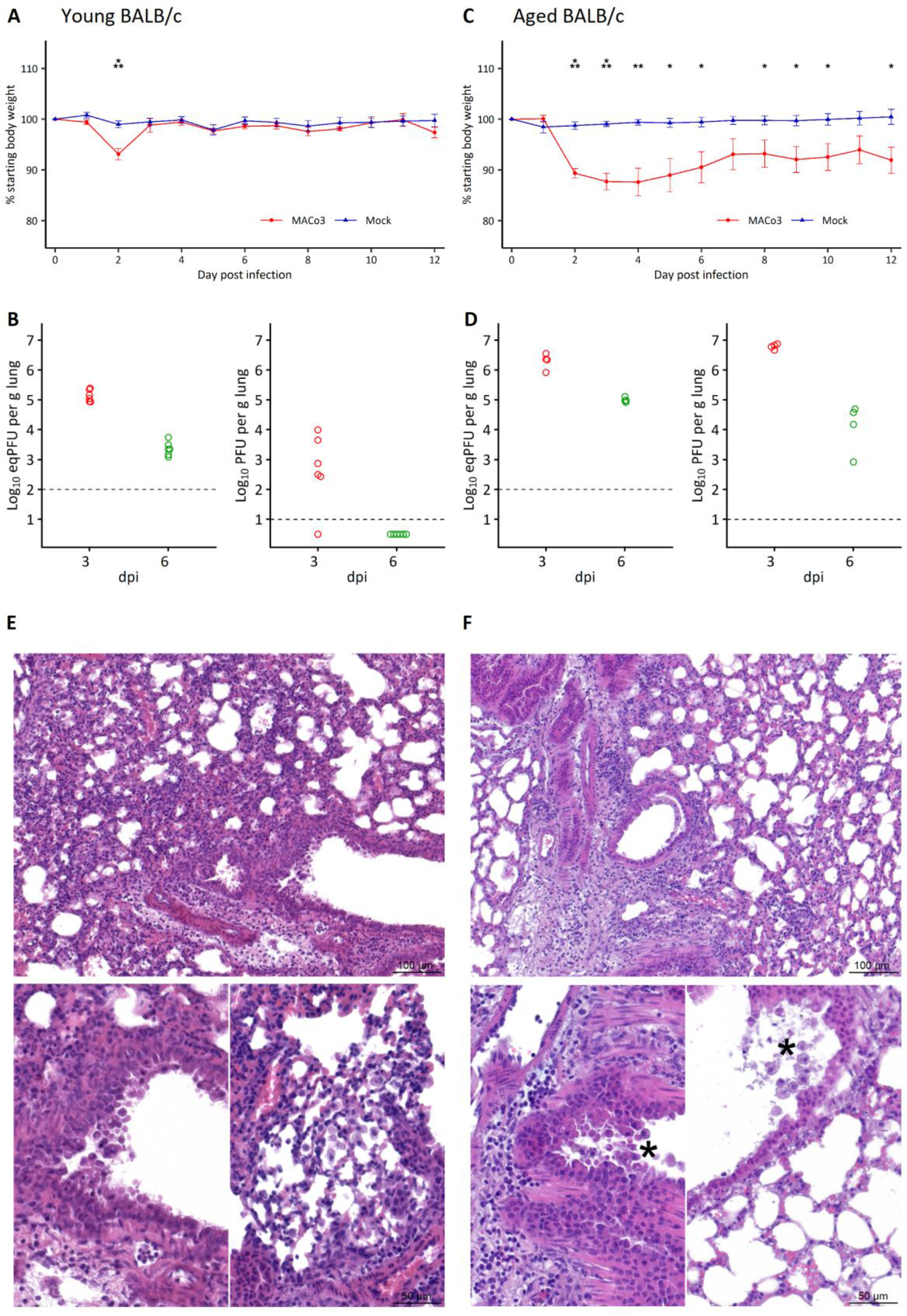
MACo3 replicates in BALB/c mouse lungs and induces age-dependent pathology. A: Body weight variations in 9-week-old BALB/c females after inoculation with PBS (n=8) or with 10^5^ PFU of MACo3 (n=8). B: Viral load and viral titer in the lung of 9-week-old BALB/c females 3 days (n=6) or 6 days (n=6) after inoculation with 10^5^ PFU of MACo3. C: Body weight variations in 53-55-week-old BALB/c males after inoculation with PBS (n=6) or with 10^5^ PFU of MACo3 (n=6). B: Viral load and viral titer in the lung of 76-90-week-old BALB/c females 3 days (n=4) or 6 days (n=4) after inoculation with 10^5^ PFU of MACo3. The dotted line represents the limit of detection (LOD), and undetected samples are plotted at half the LOD. E. H&E staining of lung section from young BALB/c mice. Top: Minimal broncho-interstitial pneumonia and perivasculitis. Bottom: Minimal alteration/necrosis of bronchiolar epithelial cells (left) and alveolar infiltrate of macrophages (right). F. Aged BALB/c mice. Top: More severe broncho-interstitial pneumonia. Bottom: Endothelitis (arrowhead) and necrosis of bronchiolar epithelial cells (star). *: p<0.05; **: p<0.01; ***: p<0.001

Infection of young C57BL/6 mice similarly led to an asymptomatic infection with moderate body weight loss peaking at dpi 2, but the difference with the mock-infected group persisted longer than in BALB/c mice. Weight decrease was more pronounced in aged C57BL/6 mice (peaking at 14% on dpi 3-4 Fig 3C) but mice developed only mild and transient symptoms (ruffled fur, hunched back posture and breathing difficulties). Viral replication in the lungs was less homogenous than in BALB/c mice, with a similar pattern of higher and longer detection of virus with increased age (Fig 3D). Like in BALB/c mice, the virus did not disseminate to other organs although low viral loads were detected in the feces of three out of four aged C57BL/6 mice at dpi 3 (Fig S2). Histopathological analysis revealed lesions similar to those observed in BALB/c mice. However, in comparison with BALB/c mice, inflammatory lesions were slightly more severe in young animals (Fig 3E), and the severity of the lesions was more heterogeneous in aged mice (Fig 3F).

**Figure 3:**
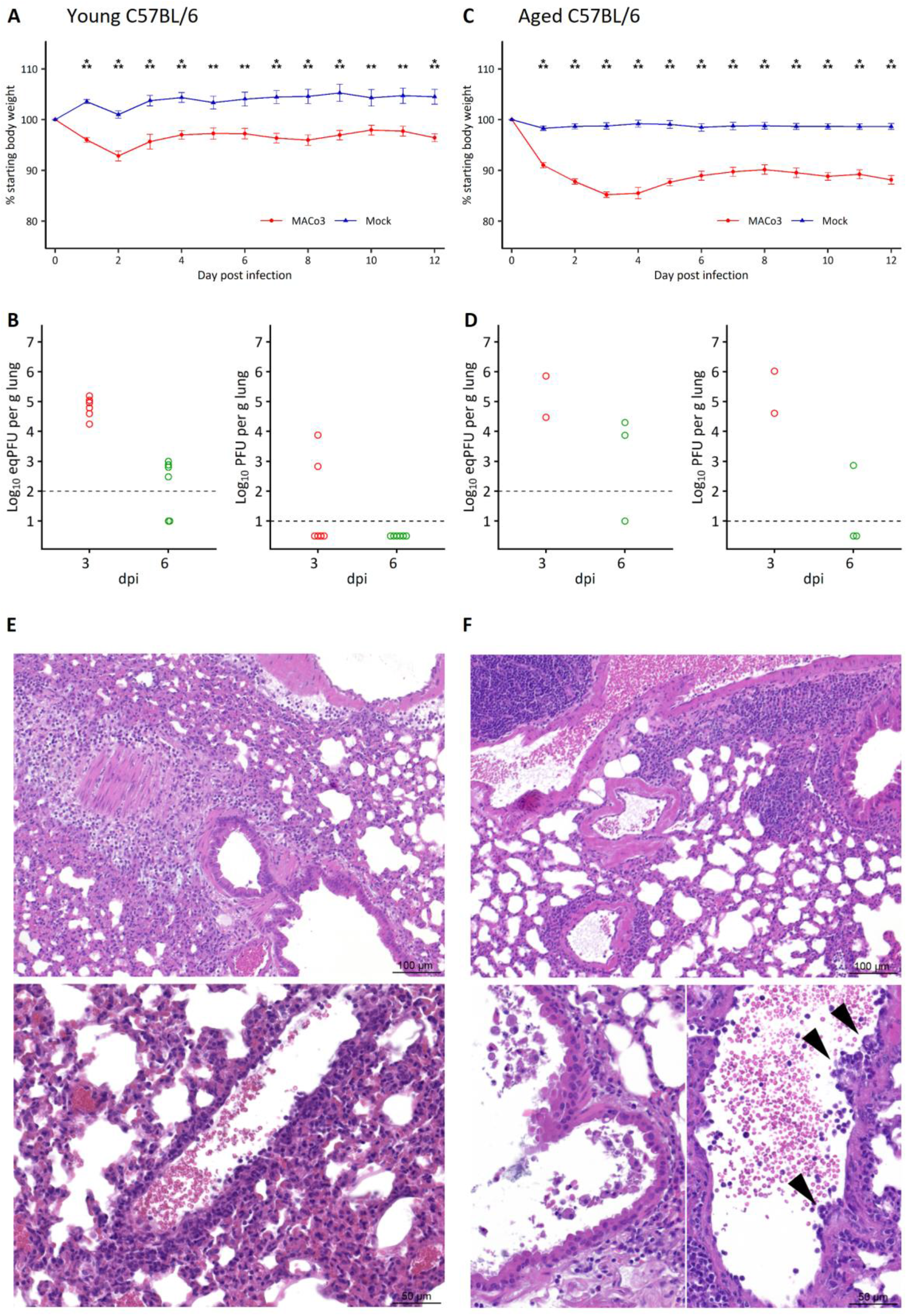
MACo3 replicates in C57BL/6 mouse lungs and induces age-dependent pathology. A: Body weight variations in 9-week-old C57BL/6 females after inoculation with PBS (n=8) or with 10^5^ PFU of MACo3 (n=8). B: Viral load and viral titer in the lung of 9-week-old C57BL/6 females 3 days (n=6) or 6 days (n=6) after inoculation with 10^5^ PFU of MACo3. C: Body weight variations in 50-week-old C57BL/6 males after inoculation with PBS (n=7) or with 10^5^ PFU of MACo3 (n=7). B: Viral load and viral titer in the lung of 58-week-old C57BL/6 males 3 days (n=2) or 6 days (n=3) after inoculation with 10^5^ PFU of MACo3. The dotted line represents the limit of detection (LOD), and undetected samples are plotted at half the LOD. E: H&E staining of lung section from young C57BL/6 mice. Top: Mild broncho-interstitial pneumonia with slightly more severe inflammation than in young BALB/c mice. Bottom: Endothelitis and perivasculitis. F: Aged C57BL/6 mice. Top: More severe broncho-interstitial pneumonia (in two out of four mice, the other mice displayed only minimal lesions). Bottom: Necrosis of bronchiolar epithelial cells (left) and endotheliitis (right, arrowheads). *: p<0.05; **: p<0.01; ***: p<0.001

MACo3 is a novel mouse-adapted SARS-CoV-2 strain which carries a small number of mutations. Its ability to replicate in mice is likely due to amino acid changes at positions 493 and 498 within the spike RBD. Changes at either or both these positions have been repeatedly detected in SARS-CoV-2 viruses passaged in mice [8,10-12], and, in addition to N501Y [9,14], appears to constitute parsimonious evolutionary paths to productive interaction with the murine version of ACE2 (Fig 1D). Concordantly, Huang and colleagues similarly reported a mouse adapted strain carrying changes at positions 493 and 498, but to other basic amino-acids (K and H respectively). Retrospective analysis revealed that the Q493R mutation was fixed in early passages (Fig 1C), enabling consistent replication in BALB/c mice (Fig 1B), prior to the fixation of Q498R. In other reports, amino acid changes at position 493 appeared secondarily, and were associated with increased pathogenicity for two SARS-CoV-2 strains: MA to MA10 [10] and MASCp6 to MASCp36 [12]. Of note, like other previously reported mouse adapted stains, MACo3 also carries changes outside of the spike which may contribute to the virus fitness and pathogenic profile in mice.

MACo2 and MACo3 expand the collection of mouse-adapted variants which are instrumental to investigate the molecular mechanisms of species-specific interactions between SARS-CoV-2 spike and the ACE2 receptor. Compared to other mouse-adapted variants, inoculation with MACo3 results in mild to moderate symptoms in the two laboratory mouse strains tested, despite high replication in the lungs. We previously reported similar observation upon infection with the beta (B.1.351) and gamma (P.1) variants of concern [14]. Overall, C57BL/6 mice were less susceptible to MACo3 inoculation than BALB/c mice, a difference which has been previously reported with another mouse-adapted SARS-CoV-2 strain [10]. Although more severe mouse models (in particular the K18-hACE2 transgenic model) have been used to test vaccine or antibody efficacy by assessing protection against mortality [15,16], body weight loss, viral replication in the respiratory tract and lung pathology are hallmarks of SARS-CoV-2 infection which are sufficient to test countermeasures efficacy [6,17]. We have successfully used MACo3 to demonstrate the protection conferred by a measles-vectored COVID-19 vaccine [18].

CC strains CC071 and CC001 were instrumental to adapting the virus to mice. They showed some permissiveness to the replication of an early isolate of SARS-CoV-2 (Pango lineage B), unlike C57BL/6 mice in which no virus was detected upon infection (Fig S1). Although others were successful, we also attempted twice to adapt a B.1 virus (carrying the D614G mutation) on aged BALB/c mice, without success. Most interestingly, CC001 has inherited its *Ace2* allele from 129S1/SvImJ, while CC071 carries a NOD/ShiLtJ allele (http://csbio.unc.edu/CCstatus/CCGenomes/). Both alleles encode the same protein sequence as the C57BL/6 allele, suggesting that *Ace2* is not responsible for the permissiveness of CC001 and CC071. As they can replicate in any mouse genetic background, mouse-adapted SARS-CoV-2 viruses such as MACo3 will be useful to identify host genetic and non-genetic factors influencing the severity of SARS-CoV-2 infections in mice.

## Acknowledgements

We are grateful to Dr Luis Enjuanes (National Center for Biotechnology, Spain) for the gift of the DBT cells expressing mACE2. We thank Magali Tichit and David Hardy (Histology platform, Institut Pasteur) and Hélène Huet and Jean-Luc Servely (Histology and Pathology Unit, EnvA) for the technical processing of histological samples and the histology pictures, and the staff of the BSL-3 mouse facility. We also thank the team of the core facility P2M (Institut Pasteur) for genomic sequencing.

## Funding

This work was supported by the « URGENCE COVID-19 » fundraising campaign of Institut Pasteur), the French Government’s Investissement d’Avenir program, Laboratoire d’Excellence Integrative Biology of Emerging Infectious Diseases (Grant No. ANR-10-LABX-62-IBEID), the Agence Nationale de la Recherche (Grant No. ANR-20-COVI-0028-01) and the RECOVER project funded by the European Union’s Horizon 2020 research and innovation programme under grant agreement No. 101003589. ESL acknowledges funding from the INCEPTION programme (Investissements d’Avenir grant ANR-16-CONV-0005).

## Author contribution

XM and ESL designed and coordinated the study. MP, LL and LC performed in vitro experiments and viral quantification. XM, LC and JJ performed in vivo experiments. GJ and ERG performed histopathological analysis. FD and MA produced the IDF0372 viral isolate under the supervision of SvdW. XM and ESL wrote and revised the manuscript with input from all authors.

## Competing interests

The authors declare no competing interests.

## Data and materials availability

All data are available in the main text or the supplementary materials. Sequence data that support the findings of this study have been deposited in the European Nucleotide Archive under study PRJEB46205.

## Supplementary Materials

### Materials and Methods

#### Ethics

All animal work was approved by the Institut Pasteur Ethics Committee (projects dap 200008 and dap 210050) and authorized by the French Ministry of Research under #24613 and #31816 in compliance with the European and French regulations on the protection of live vertebrates.

#### Cells and viruses

VeroE6 cells (African green monkey kidney cells from ATCC (CCL-81)) and DBT-mACE2 cells (a kind gift of Dr Luis Enjuanes) were maintained in Dulbecco’s Modified Eagle’s Medium (DMEM) supplemented with 10% fetal bovine serum (FBS) and penicillin/streptomycin. DBT-mACE2 cells were supplemented with 800 μg/ml of G418 to maintain the plasmid expressing murine ACE2.

The B (Pango lineage) strain: hCoV-19/France/IDF-0372-isl/2020, GISAID accession Id: EPI_ISL_410720; and B.1 strain: hCoV-19/France/GES-1973/2020, GISAID accession Id: EPI_ISL_414631; were supplied by the National Reference Centre for Respiratory Viruses hosted by Institut Pasteur (Paris, France) and headed by Pr. S. van der Werf. The human sample from which the IDF-0372 strain was isolated has been provided by X. Lescure and Y. Yazdanpanah from the Bichat Hospital. Viruses were amplified and titrated by standard plaque forming assay on VeroE6 cells. The sequence of the stocks was verified by RNAseq on the mutualized platform for microbiology (P2M). All work with infectious virus was performed in biosafety level 3 containment laboratories at Institut Pasteur.

#### Mouse strains

BALB/cJRj (BALB/c) and C57BL/6JRj (C57BL/6) mice were purchased from Janvier Labs (Le Genest St Isle, France). Collaborative Cross mice were purchased from the Systems Genetics Core Facility, University of North Carolina, and bred at the Institut Pasteur. Mice were maintained under specific-pathogen-free conditions with a 14-h light and 10-h dark cycle and ad libitum food and water in the Institut Pasteur animal facility.

#### *In vivo* passaging and infection

Infection studies were performed in animal biosafety level 3 (BSL-3) facilities at the Institut Pasteur, in Paris. To initiate serial passages, 10^5^ PFU of IDF0372 were first inoculated to two anesthetized (ketamine/xylazine) 17-week-old CC071 males which were euthanized by ketamine/xylazine overdose at dpi 3 to collect their lungs. Lung homogenate was prepared from each mouse as previously described [14]. Twenty microliters of a 1:1 mix of the two homogenates were immediately inoculated to two other CC071 mice. The viral load was determined in each homogenate by RT-qPCR. The procedure was repeated on CC071, CC001 and C57BL/6 mice as described on Fig 1A. Lungs were collected at dpi 2 or 3. At every passage, mice received a mix of lung homogenates from all mice of the previous passage. The populations of viruses present in the homogenates at P15 and P16 were cultured, identified as MACo1 and MACo2, tittered and sequenced. Further mouse adaptation was performed by passaging on 6-18 month-old BALB/c mice. The first pair (P17) was inoculated with 4.4×10^4^ PFUs of MACo2. Passages were performed as described above until P27 from which viruses were isolated and cloned. One of the clones, called MACo3, was amplified and sequenced. This clone was used for all infection experiments.

Infection experiments on BALB/c and C57BL/6 mice were performed as above with 10^5^ PFU of MACo3 (sex, age and number of mice are indicated for each mouse group in the figure legends). Clinical signs of disease and weight loss were monitored daily. Mice were euthanized by ketamine/xylazine overdose at the end of the observation period (dpi 12) or at indicated time points (dpi 3 or 6). Samples for viral load (right lung lobe, internal organs and feces) and histopathological analyses (left lung lobe) were collected, immediately frozen in dry ice and stored at −80°C. The left lung lobe was fixed by submersion in 10% phosphate buffered formalin for 7 days prior to removal from the BSL3 for processing. The right lung lobe was placed on a 70μ cell strainer (Falcon), minced with fine scissors and ground with a syringe plunger using 400 μl of PBS. Lung homogenates were used for virus quantification.

#### Virus quantification

Tissue samples (brain, liver, spleen, heart, jejunum) and feces collected in the colon were weighed and homogenized in PBS using Lysing Matrix D tubes (MP biomedicals) and MP biomedicals tissue homogenizer. After centrifugation, RNA was extracted from 200 μl of lysates using NucleoMag Pathogen kit (Macherey Nagel). Viral RNA quantification was performed by quantitative reverse transcription PCR (RT-qPCR) using the IP4 set of primers and probe as described on the WHO website (https://www.who.int/docs/default-34source/coronaviruse/real-time-rt-pcr-assays-for-the-detection-of-sars-cov-2-institut-35pasteur-paris.pdf?sfvrsn=3662fcb6_2) and the Luna Universal Probe One-Step RT-qPCR Kit (NEB).

For plaque assay, 10-fold serial dilutions of samples in DMEM were added onto VeroE6 monolayers in 24 well plates. After one-hour incubation at 37°C, the inoculum was replaced with 2% FBS DMEM and 1% Carboxymethylcellulose. Three days later, cells were fixed with 4% formaldehyde, followed by staining with 1% crystal violet to visualize the plaques.

#### Histological analysis

Histological analysis was performed on paraffin-embedded 4μm-thick sections used for hematoxylin-eosin (H&E) staining.

#### Statistical analysis

Body weight loss was compared between control and MACo-3-infected mice by one-way ANOVA. Data were analyzed and graphed using R software.

**Figure S1:**
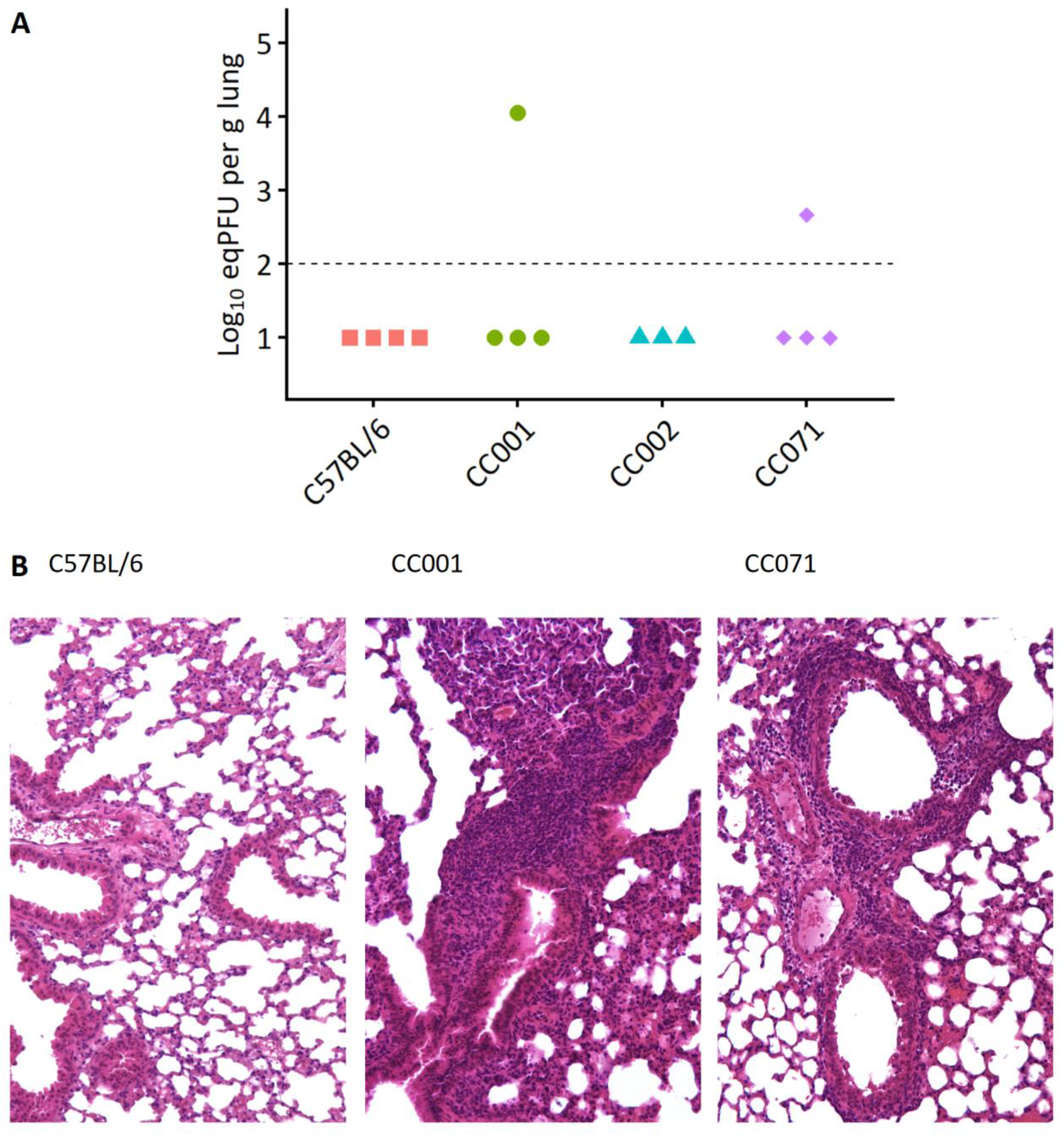
Viral replication and lung lesions after infection of C57BL/6 and three CC strains with wildtype IDF0372 SARS-CoV-2. A: Viral load in the lungs of 9-16-week-old C57BL/6, CC001, CC002 and CC071 males inoculated with 4×10^4^ PFUs of IDF0372. The dotted line represents the limit of detection (LOD), and undetected samples are plotted at half the LOD. B: H&E staining of lung sections from one of the C57BL/6 (virus-negative) mice (left, with no significant histological lesions) and the virus-positive CC001 (center) and CC071 (right) mice which both exhibit broncho-interstitial pneumonia.

**Figure S2:**
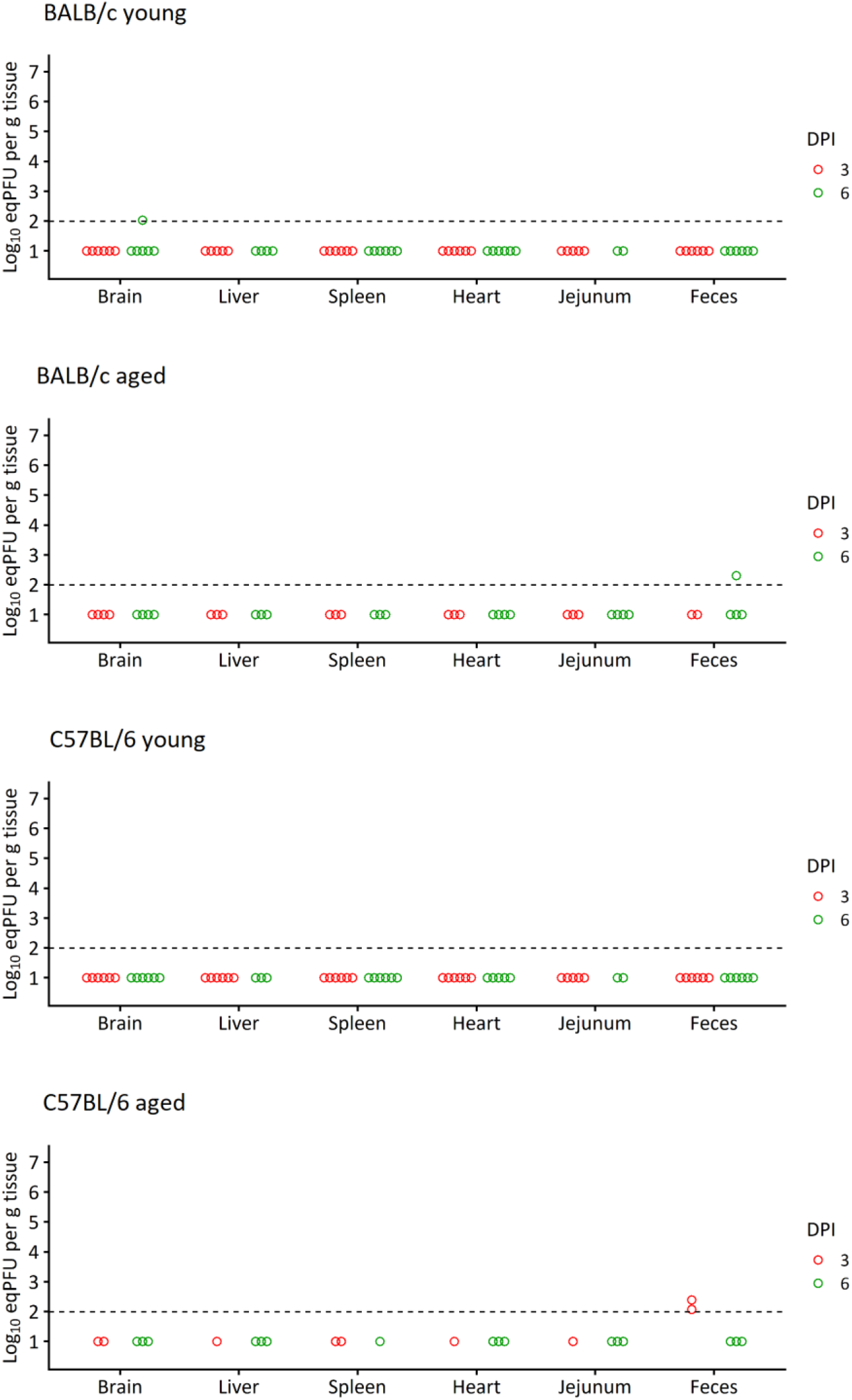
Viral loads in samples from young and aged C57BL/6 and BALB/c mice. Samples were collected on dpi 3 or 6, on the same mice as in Figures 2B, 2D, 3B and 3D, respectively. Viral load (eqPFU per g of tissue) was determined by qRT-PCR. The dotted line represents the limit of detection (LOD), and undetected samples are plotted at half the LOD.

## Notes

### Competing Interest Statement

The authors have declared no competing interest.

